# Applicability of epigenetic age models to next-generation methylation arrays

**DOI:** 10.1101/2024.06.07.597709

**Authors:** Leonardo D. Garma, Miguel Quintela-Fandino

## Abstract

**Background:** Epigenetic clocks based on DNA methylation data are routinely used to obtain surrogate measures of biological age and estimate epigenetic age acceleration rates. These tools are mathematical models that rely on the methylation state of specific sets of CpG islands quantified using microarrays. The set of CpG islands probed in the microarrays differed between the models. Thus, as new methylation microarrays are developed and older models are discontinued, existing epigenetic clocks might become obsolete. Here, we explored the effects of the changes introduced in the new DNA methylation array from Illumina (EPICv2) on existing epigenetic clocks.

**Methods:** We compiled a whole-blood DNA methylation dataset of 10835 samples to test the performance of four epigenetic clocks on the probe set of the EPICv2 array. We then used the same data to train a new epigenetic age prediction model compatible across the 450k, EPICv1 and EPICv2 microarrays. We compiled a validation dataset of 2095 samples to compare our model with a state-of-the-art epigenetic clock. Using two datasets with repeated samples from the same subjects, we computed an estimate of the contribution of technical noise and intra-subject variation to the variation of epigenetic age predictions from each of the models tested. We used a dataset of cancer survivors who had undergone different types of therapy, a dataset of breast cancer patients and controls, and a dataset from an exercise-based interventional study to test the ability of our model to detect alterations in epigenetic age acceleration.

**Results:** We found that the results of the four epigenetic clocks tested are significantly distorted by the absence of specific probes in the EPICv2 microarray, causing an average difference of up to 25 years. We developed an epigenetic age prediction model compatible with the 450k, EPICv1 and EPICv2 microarrays. Our model produced highly accurate chronological age predictions that were comparable to those of a state-of-the-art epiclock. We obtained estimates for the variation of epigenetic age acceleration on normal, non-pathological populations associated with each of the models tested. These parameters provide thresholds to evaluate the relevance of epigenetic age alterations. In all cases, the estimated technical noise and intra-subject variability were smaller than the population-based epigenetic age prediction variability. Finally, we used our new models to reproduce previous results showing increased epigenetic age acceleration in cancer patients and in survivors who had been treated with radiation therapy, as well as a lack of changes as a result of exercise-based interventions.

**Conclusion:** Our work demonstrated that existing epigenetic clocks need to be updated to be applicable to data generated with the new EPICv2 microarray, which has phased out the 450k and EPICv1 models. To overcome this technical hurdle, we developed a new model that translates the capabilities of state-of-the-art epigenetic clocks to the new EPICv2 platform and is cross-compatible with the 450k and EPICv1 microarrays. Our characterization of the variation of epigenetic age predictions provides useful metrics to contextualize the biological relevance of epigenetic age alterations. The analysis of data from subjects influenced by radiation, cancer and exercise-based interventions shows that despite being good predictors of chronological age, neither a pathological state like breast cancer, a hazardous environmental factor (radiation) or exercise (a beneficial intervention) caused significant changes in the values of the “epigenetic age” determined by these first-generation models.

## Introduction

In the rapidly evolving field of epigenomics, the development of epigenetic clocks has revolutionized our ability to gauge biological aging through DNA methylation patterns. Changes in the methylation state of CpG sites have proven to be highly correlated with chronological age^1,2^. Thus, DNA methylation patterns are being increasingly used to gain insights into aging and associated pathologies^2^ and have led to the development of several epigenetic clocks, which are predictive models that estimate the age of subjects based on methylation markers^3^. In these epigenetic clocks, the model prediction is interpreted as the “biological” or “epigenetic” age. Subjects whose biological age is greater than their chronological age are considered to have epigenetic age acceleration^4^ (EAA). EAAs have been linked to increased risk for various health issues and mortality, independent of traditional risk factors^5–8^. Epigenetic clocks have thus not only underscored the potential of DNA methylation as a biomarker of aging^9^ but also represent a valuable tool to gain insight into the complexities of multiple pathologies^4^. In cancer, EAAs have been linked to an increased risk of developing breast and colon cancer^10,11^. Moreover, breast tissue from breast cancer patients has been shown to exhibit EAA^12^; likewise, cancer survivors do present an accelerated epigenetic age compared to noncancer patients, with an acceleration rate dependent on the treatment intensity received^13^.

From the initial model of Bockland *et al*. published in 2011^1^, the field quickly evolved toward more robust models, such as the Hannum^14^ and the Horvath^15^ clocks, and later toward “second generation” approaches, which included features beyond DNA methylation with the promise of providing a “biological age” prediction that could better reflect age-related health status^16–18^. The field continues to evolve, with the implementation of new predictive models^19–21^ and the development of species-^22,23^ and tissue-specific^24–27^ epigenetic clocks. Most recently, Bernabeu *et al*. conducted a large-scale study to refine the predictive ability of both first (methylation-based) and second (including other features) generation epigenetic clocks, which yielded a significant improvement over existing models in the ability to predict chronological age from DNA methylation data across multiple cohorts with subjects of all ages^28^.

The evolution of epigenetic clocks has run in parallel with that of methylation microarrays, whose coverage of DNA methylation sites increased from 27578 in 2008^29,30^ to 866836 in 2015^31^. In 2023, Illumina released the newest iteration of its series of methylation arrays, EPICv2, which targets 936866 methylation sites^32^. Although the vast majority of probes have been conserved across different microarray models over the years^32–34^, some of them have been lost. The probes on the latest (and currently the only commercially available) methylation microarray version cover more than 80% of the probes used by the most popular epigenetic clocks (PhenoAge^16^, Horvath (2013)^15^, Horvath (2018)^35^, Hannum^14^, DunedinPACE^17^, etc.), but they do not offer complete coverage for any of them^36^. Given that earlier microarray models are now discontinued while there is still a need to use epigenetic clocks, it is necessary to evaluate whether the existing models can offer reliable results when using EPICv2 data.

Recent studies have shown that DNA methylation is a dynamic molecular feature, exhibiting changes which can affect epigenetic age predictions even within a 24-hour timeframe^37–39^. Apsley et al. demonstrated that methylation values of probes in the EPICv1 array do exhibit significant changes using repeated sampling on the same subjects on 4 different timepoints within a 5-hour interval. The effect was observed on multiple probes used by common epigenetic clocks, although the changes in the epigenetic age predictions were not directly reported^38^. More recently, Koncevičius et al. reported oscillations in the epigenetic age predictions produced by 17 epigenetic age models on 48 samples from the same subject obtained during a 72-hour period, which they attributed to circadian oscillations^37^. Therefore, it is necessary to quantify the range of spontaneous variation in epigenetic age predictions to be able to distinguish truly biologically relevant alterations.

In this study, we examined the performance of existing epigenetic clocks using the CpG probes available in the new EPICv2 methylation array in human blood samples. We observed that the biological age predictions of these models were distorted due to missing probes in the EPICv2 array. To overcome this hurdle, we trained a model that can be applied to data obtained with 450k, EPICv1 or EPICv2 chips and whose performance is at least on par with that of state-of-the-art models. We then estimated the variation of EAA in normal, non-pathological populations for our model and for four existing epiclocks. We used two datasets with repeated samples from the same subjects to obtain estimates of the contributions of technical and biological, intra-subject noise to the variation observed in the EAA distributions. The extent of these variations can set a threshold to distinguish normal from biologically relevant EAA changes. Finally, our model reproduced previously reported EAA differences attributed to radiation therapy and to sporadic breast cancer, and the negative results from an interventional study using physical exercise.

## Results

### DNA methylation datasets

We explored public repositories to assemble a large and diverse DNA methylation dataset to examine the performance of existing epigenetic clocks and to train and test new models. We used public data to generate two separate datasets: 1) a main dataset for evaluating existing models and training new ones and 2) a validation dataset to test the generalizability of new models.

The main dataset included whole blood DNA methylation data of human subjects from 24 previous studies (Supplementary Table 1). We filtered the data to retain exclusively subjects labeled as controls, with reported age and sex values, resulting in a total of 11825 individuals. To be able to split the data in a stratified fashion into training and test sets, we kept only the samples with an age value present at least twice in the dataset, leaving a total of 10835 subjects (Figure 1A). The subjects in this main dataset had ages between 8 and 96 years, with a mean of 47.68 years and an almost even distribution of sexes (5456 females and 5379 males) (Figure 1B).

**Figure 1.**
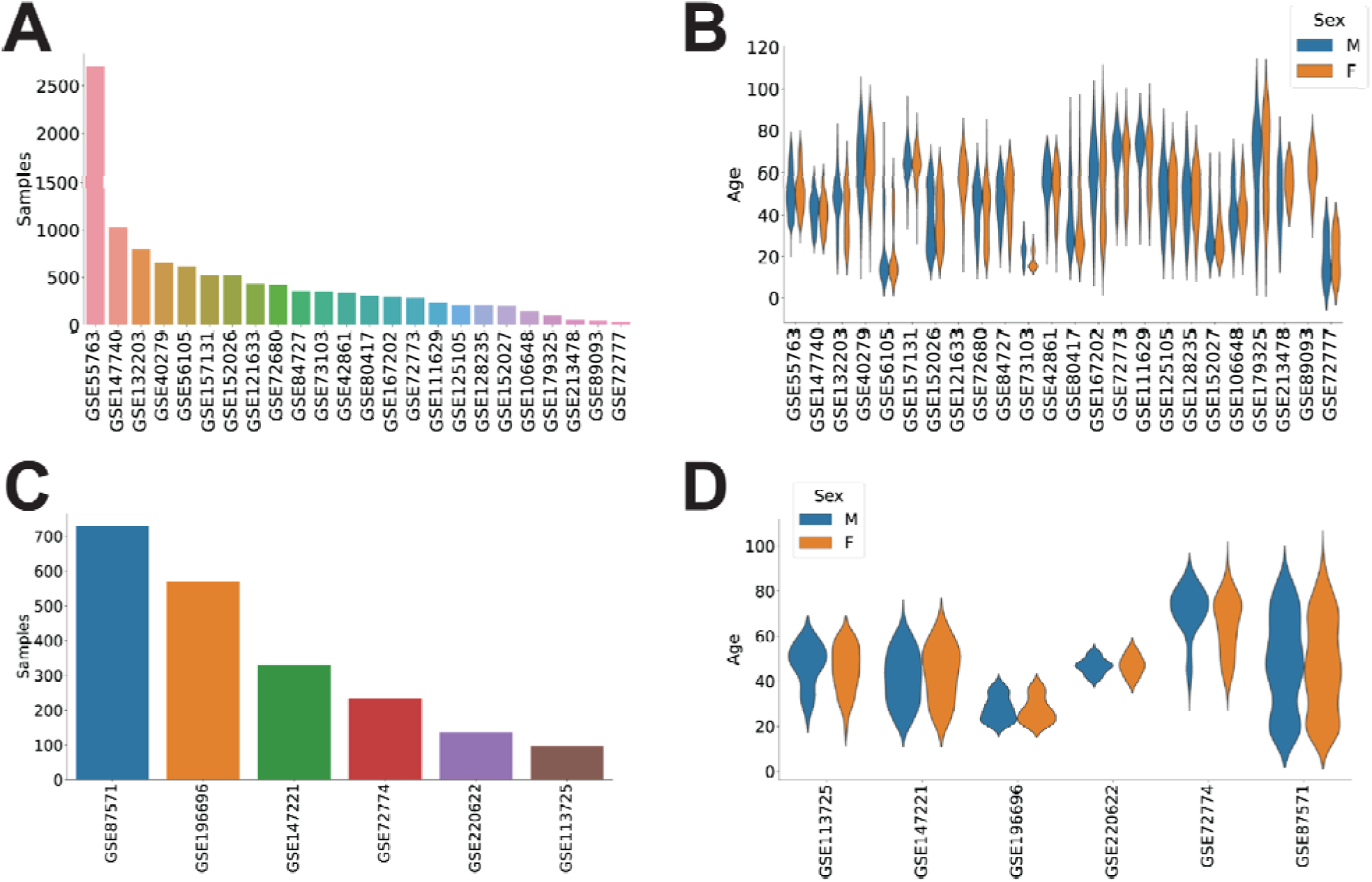
Public DNA methylation datasets compiled for the study. **A**. Number of subjects present in each of the datasets included in the main set. **B.** Distribution of ages and sex in the datasets included in the main set. **C** and **D**. Distribution of the number of subjects and their age and sex in the datasets used for the validation set.

The validation dataset was composed of data from 6 additional studies (Supplementary Table 1). We selected only control subjects with reported age and sex values, which resulted in a total of 2095 individuals. The age range in this validation set was between 14 and 94 years, and the sex distribution was slightly biased, with 828 females and 1267 males (Figure 1C-D).

### Existing epigenetic clocks generate distorted results from EPICv2 data

Although more than 80% of the probes used by existing epiclocks are conserved in the EPICv2 methylation array^36^, applying these models to data limited to the probes present in the EPICv2 array results in differences in epigenetic age predictions. We applied four different models (Horvath^15^, Hannum^14^, PhenoAge^16^ and the recent chronological age prediction model (cAge) from Bernabeu *et al*.^28^) to the main dataset of 10835 individuals, using either all the required probes (*complete* models) or only those present in the new EPICv2 array (*truncated* models). The proportion of probes relevant for these models in the EPICv2 ranged from 84.5% (Hannum) to 97.1% (cAge).

We observed a high degree of correlation between chronological age and the predicted biological age in all cases, with cAge obtaining the highest value (r=0.979). After the models were truncated to the EPICv2 probeset, they maintained this correlation, which was still strong (r>0.88) in all cases (Figure 2A).

**Figure 2.**
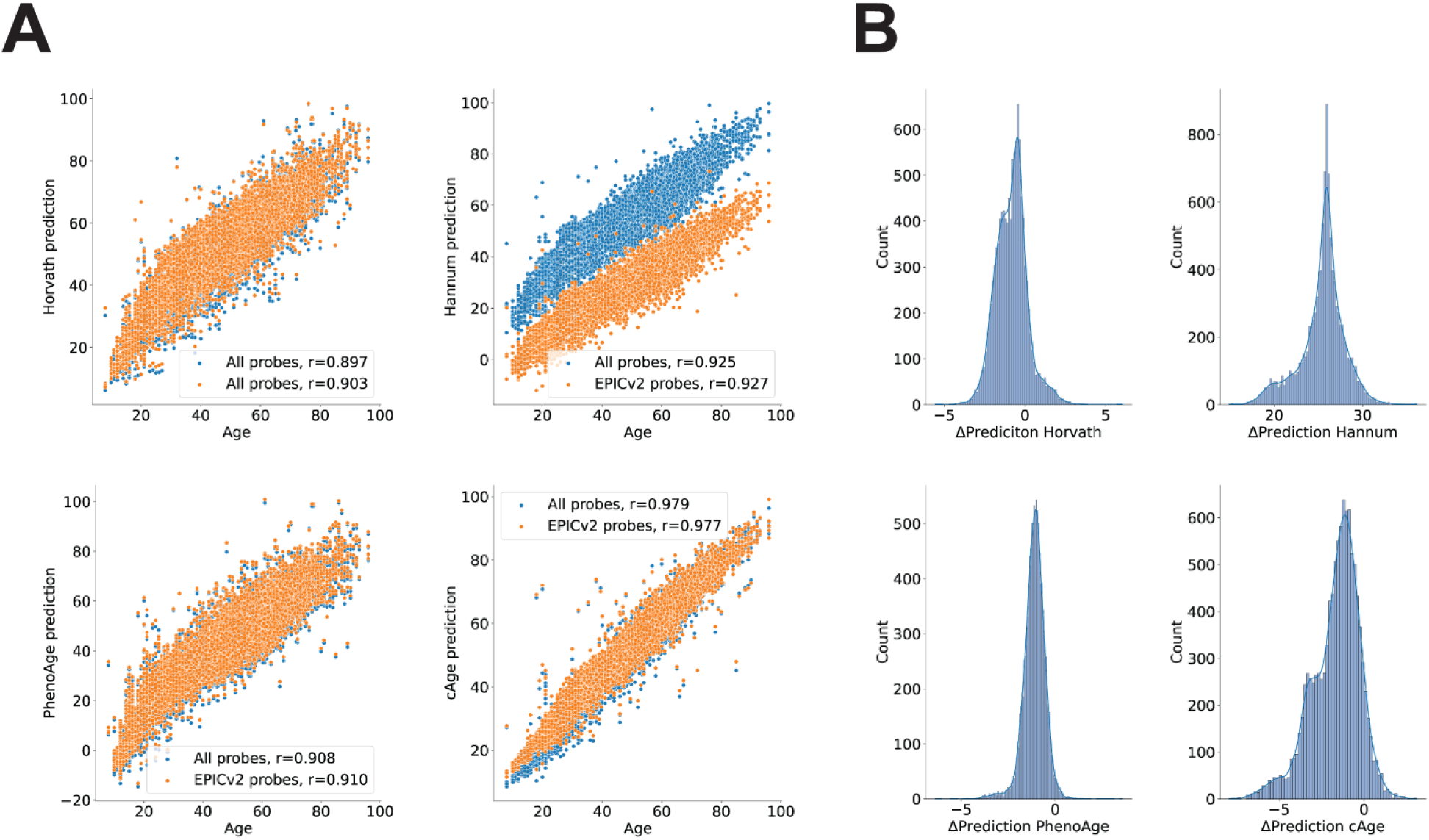
Epigenetic clock age prediction drifts caused by the loss of probes in the EPICv2 microarray. **A**. Epigenetic age predictions by four existing epigenetic clocks in 10835 subjects using either their complete sets of CpG probes (blue) or only those available in the EPICv2 array (orange). **B**. Distribution of differences between the values predicted using the full sets of probes or only those present in the EPICv2 array (drift) for the same four epigenetic clocks.

However, limiting the number of probes distorted the epigenetic age predictions in all the models. In all four epiclocks, we observed a significant difference (p<0.001, one sample t-test) in the biological age predicted by the complete and the truncated models (Figure 2B). This effect was greater on the Hannum clock model, with an average difference of 25.45 years. The larger models PhenoAge (513 parameters) and cAge (4058 parameters) had average differences of −1.11 and −1.67 years respectively, whereas the average difference on the Horvath clock was just −0.82 years.

To characterize the performance of these models in predicting chronological age, we computed the Pearson correlation coefficient between the predictions and chronological age (r), the mean squared error (MSE) and median absolute error (MAE) for each of them. We computed the mean (μ_EAA_) and standard deviations (σ_EAA_) of the EAA distributions associated to each epigenetic (Table 1).

**Table 1.** Metrics from all the complete (C) and truncated (T) models on the different datasets used in the study.

|  |  | Hannum<sub>C</sub> | Hannum<sub>T</sub> | Horvath<sub>C</sub> | Horvath<sub>T</sub> | PhenoAge<sub>C</sub> | PhenoAge<sub>T</sub> | cAge<sub>C</sub> | cAge<sub>T</sub> | General Model | Combined Model |
| --- | --- | --- | --- | --- | --- | --- | --- | --- | --- | --- | --- |
| Main dataset | **r** | 0,92 | 0,93 | 0,90 | 0,90 | 0,91 | 0,91 | 0,98 | 0,98 | N/A | N/A |
|  | **MAE (years)** | 7,58 | 19,08 | 6,37 | 6,65 | 7,05 | 6,44 | 2,55 | 3,57 | N/A | N/A |
|  | **MSE (years)** | 81,58 | 410,80 | 65,85 | 69,57 | 78,43 | 66,92 | 12,82 | 20,88 | N/A | N/A |
|  | **μ<sub>EAA</sub> (years)** | 6,38 | -19,06 | 3,32 | 4,14 | -5,25 | -4,14 | 1,07 | 2,73 | N/A | N/A |
|  | **σ<sub>EAA</sub> (years)** | 6,39 | 6,89 | 7,40 | 7,24 | 7,13 | 7,06 | 3,42 | 3,67 | N/A | N/A |
| Validation dataset | **r** | 0,96 | 0,97 | 0,94 | 0,94 | 0,94 | 0,94 | 0,99 | 0,99 | 0,99 | 0,99 |
|  | **MAE (years)** | 7,04 | 18,93 | 6,10 | 6,59 | 7,90 | 7,16 | 1,94 | 2,21 | 2,08 | 1,74 |
|  | **MSE (years)** | 68,45 | 397,10 | 55,64 | 63,44 | 89,60 | 74,98 | 6,41 | 8,32 | 7,55 | 5,96 |
|  | **μ<sub>EAA</sub> (years)** | 6,21 | -18,92 | 3,08 | 4,24 | -6,85 | -5,86 | 0,01 | 1,37 | -0,54 | -0,21 |
|  | **σ<sub>EAA</sub> (years)** | 5,47 | 6,27 | 6,80 | 6,74 | 6,53 | 6,38 | 2,53 | 2,54 | 2,70 | 2,43 |
| Koncevičius et al. | **σ<sub>noise</sub> (years)** | 1,18 | 1,01 | 1,26 | 1,19 | 1,13 | 1,09 | 0,78 | 0,86 | 0,46 | 0,50 |
|  | **σ<sub>subject</sub> (years)** | 2,20 | 1,74 | 1,99 | 1,85 | 2,37 | 2,31 | 0,98 | 1,04 | 0,89 | 0,83 |
| Apsley et al. | **σ<sub>subject</sub> (years)** | 2,06 | 1,77 | 3,31 | 3,28 | 2,61 | 2,58 | 0,99 | 1,05 | 1,30 | 0,88 |

### A new epigenetic clock compatible with the EPICv2 array

The changes in the CpG probes present in the EPICv2 array produce significant alterations in the results of existing epigenetic clocks. To overcome this limitation and to produce robust biological age predictions using EPICv2 data, we trained a new epiclock model using CpG probes common to the 450k, EPICv1 and EPICv2 methylation arrays. Based on the epigenome-wide association study (EWAS) from Bernabeu *et al*. ^28^, there are 42728 probes with a significant (p-adjusted<0.05) linear association with chronological age that are present in the three different methylation arrays. Similarly, according to the EWAS, there are 63324 CpG probes with significant (p-adjusted<0.05) quadratic associations with age which are present in the three microarray models (Figure 3A).

**Figure 3.**
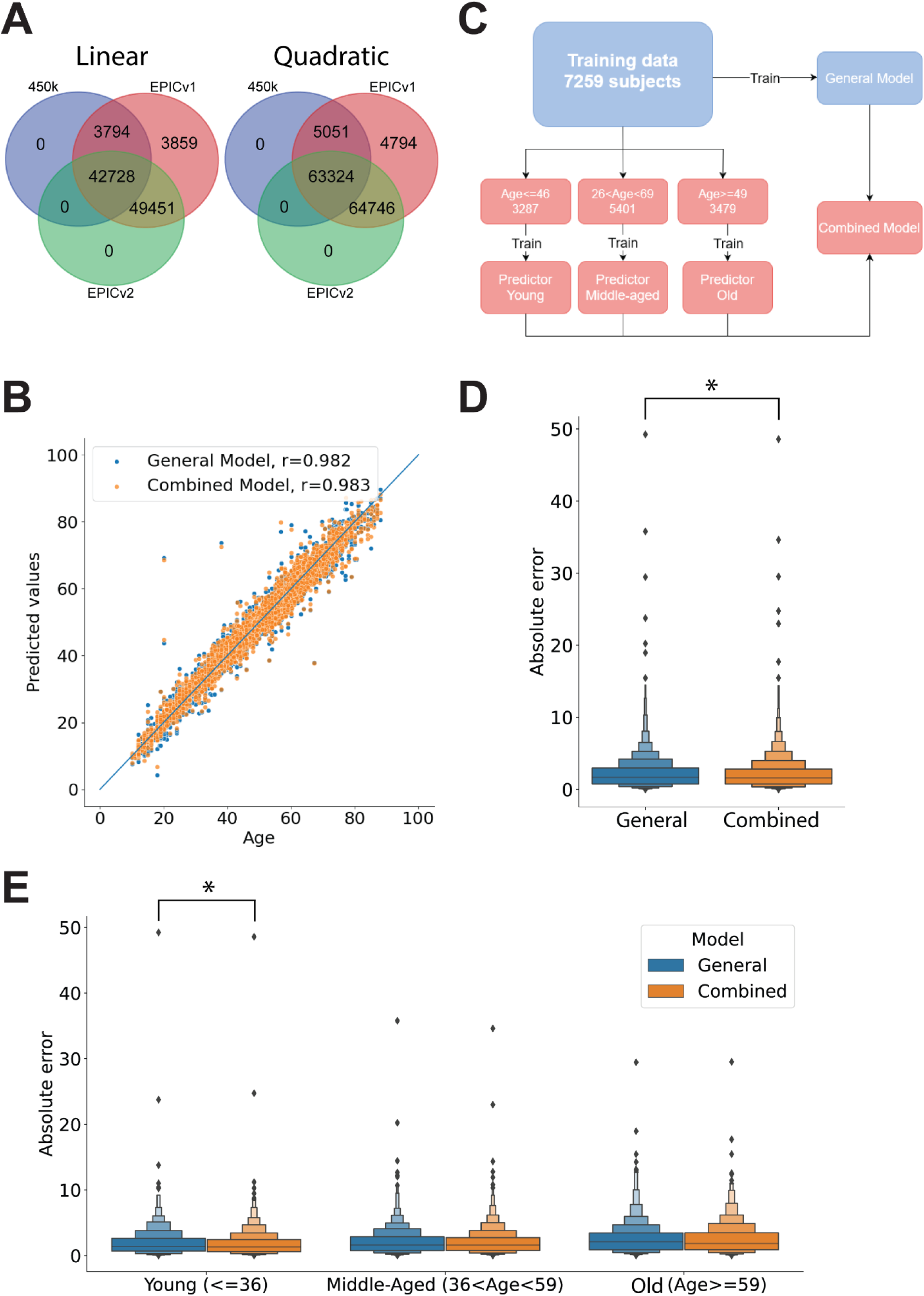
Results of the new epigenetic clocks in the test dataset. **A.** Methylation probes with significant linear (left) and quadratic (right) associations with chronological age in the EWAS from Bernabeu *et al*. present in the 450k, EPICv1 and EPICv2 arrays. **B**. Prediction results on the test set by the General and Combined models. The blue line indicates the 1:1 correspondence. **C**. Schematic overview of the training of the different age prediction models. **D**. Absolute error of the ages predicted in the test set by the general and combined models. **E**. Absolute error from the predictions on the test set by each model, broken down by age group. *Indicates significant differences (p<0.05, one-sided Wilcoxon rank-sum test) between groups.

Following the strategy employed by Bernabeu *et al*.^28^, we ranked the probes by their p-value on the EWAS and preselected the first 10000 with linear and the first 300 with quadratic associations to chronological age to train an epiclock using elastic net regression^40^ (Supplementary Table 2).

We split the main dataset with 10835 subjects into training and test sets using a 70/30 split, resulting in 7259 and 3576 samples, respectively, and trained a general age-prediction model. The model used an L1 ratio of 0.5^28^, and the alpha value was optimized to 0.001 through 5-fold cross-validation. This general model obtained a low prediction error (mean squared error (MSE)=3.14 years) and a high correlation between the predicted and chronological ages (r=0.982) in the test set (Figure 3B).

Previous works have suggested that the relationship between methylation state and age is nonlinear^14,28,41–44^. Therefore, we decided to stratify our training data into three age groups and train separate predictors for “young”, “middle-aged”, and “old” subjects. The predictors were intended to work on nonoverlapping age groups, but we did use overlapping age ranges during training to limit the inconsistencies between models assigned to contiguous groups. We produced a combined model in which the general model assigned a sample to one of the three age groups, and then the corresponding age-stratified model generated the age prediction (Figure 3C). This combined model reduced the age prediction error significantly (p<0.05, one-sided Wilcoxon rank-sum test) with respect to the general model (Figure 3B,D), lowering the MSE to 3.06 years in the test set and increasing the correlation between the predicted and chronological ages to 0.983. Although the combined model reduced the error in all age groups, the difference was statistically significant (p<0.05, one-sided Wilcoxon rank-sum test) only in the younger subjects (Figure 3E). Weights for the general and age-specific models are provided in Supplementary Table 3. The overlap of EPICv2 probes used by our general and combined models and by the rest of the epiclocks examined is presented schematically in Supplementary Figure 1A and B. To underscore the numerical differences between models, we present the coefficients used for the overlapping probes in our models, cAge and the Horvath clock in Supplementary Figure 1C.

### The performance of the new clock is on par with that of a state-of-the-art model

To examine the performance of our model, we compared it against cAge, a refined model that has shown significant improvement over previous epigenetic clocks^28^. We tested the complete cAge and our general and combined models on a validation dataset with 2095 samples from 6 studies. We included all the probes used for cAge and those required by our models and predicted the age of each sample based on the methylation data. In all cases, the predicted ages were strongly correlated with the chronological ages (Figure 4A).

**Figure 4.**
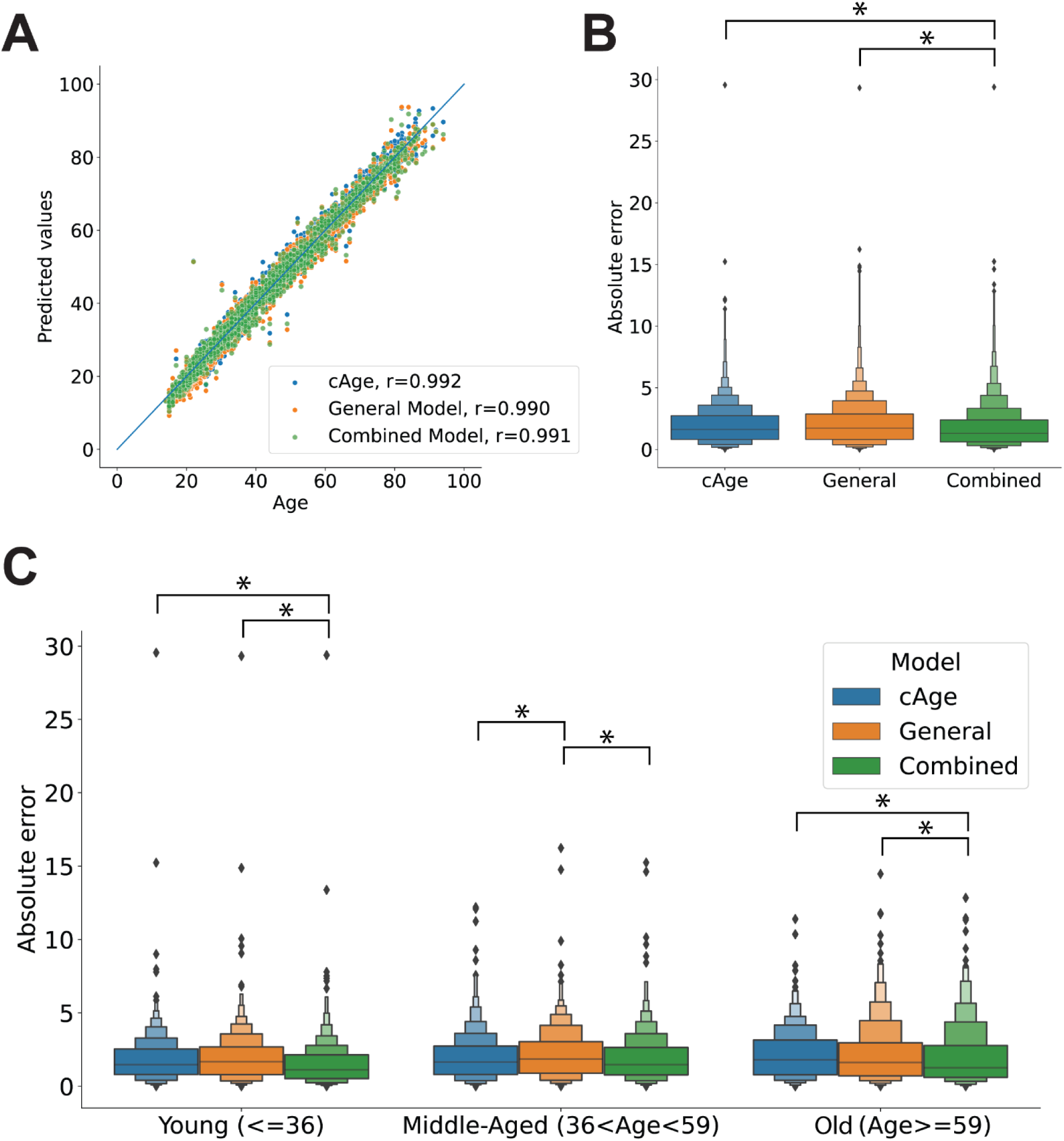
Age predictions on the validation set. **A**. Chronological age and age values predicted by cAge and the general and combined models in the validation set. **B**. Distribution of the absolute prediction error for each of the three models. **C**. Distribution of the absolute error of each model in the validation set broken down by age group. *Indicates significant differences (p<0.05, one-sided Wilcoxon rank-sum test) between groups.

Our general model performed as well as cAge, with no significant difference (p>0.05, one-sided Wilcoxon rank-sum test) in the distribution of errors between the two, whereas the error from the predictions of our combined (age-segregated) model was significantly lower (p>0.05, one-sided Wilcoxon test) (Figure 4B). By breaking down the data by age group, we could observe differences in the prediction error. In the young (<=36) and old (>=59) groups, the error from the combined model was significantly lower (p>0.05, one-sided Wilcoxon test) than that of the cAge and the general models. In the group of middle-aged subjects, the error of the general model was significantly greater (p>0.05, one-sided Wilcoxon test) than that of the other two models (Figure 4C).

To be able to perform a more general comparison, we computed again the values of r, MSE, MAE, μ_EAA_ and σ_EAA_ for our models and of the other four epigenetic clocks using the validation dataset (Table 1, Supplementary Figure 2). These metrics were largely compatible with those obtained on the main dataset, with the exception of cAge, which had a large decrease in MAE, MSE, μ_EAA_ and σ_EAA_.

### Technical and intra-subject EAA variations

It has recently been shown that epigenetic clocks are affected by spontaneous variation in DNA methylation values, leading to changes in epigenetic age predictions^37,38^. Additionally, the experimental process to obtain methylation values might introduce another source of variation. In order to establish the magnitude of these variations, we applied our model and the four epiclocks discussed above to data obtained from repeated sampling of the same subjects.

First, we examined the data from the study of Koncevičius et al., which reported DNA methylation of blood samples of a 52-year-old male subject taken every 3 hours over a period of 72 hours^37^ (GSE247197). This dataset included technical replicates, which we used to estimate the EAA deviations associated with technical noise, i.e. variations across multiple experimentally determined beta values on the same sample. All the (complete) models tested exhibited variation across technical replicates (Figure 5A). To characterize this variation, we subtracted the mean value obtained on each set of replicates and then computed the standard deviation across all the mean-centered values. We interpreted the resulting measure as an estimate of the technical noise (σ_noise_). The predictions of our general model had the lowest σ_noise_ (0.78 years), whereas Horvath’s model had the highest with, reaching 1.26 years (Figure 5B, Table 1). We then computed the standard deviation across all samples in the dataset as a first estimate for the intra-subject variability (σ_subject_). The values of σ_subject_ were in all cases larger than σ_noise_, and in this case the largest and lowest values corresponded to our combined model (0.83 years) and to PhenoAge (2.36 years) respectively (Figure 5B, Table 1).

**Figure 5.**
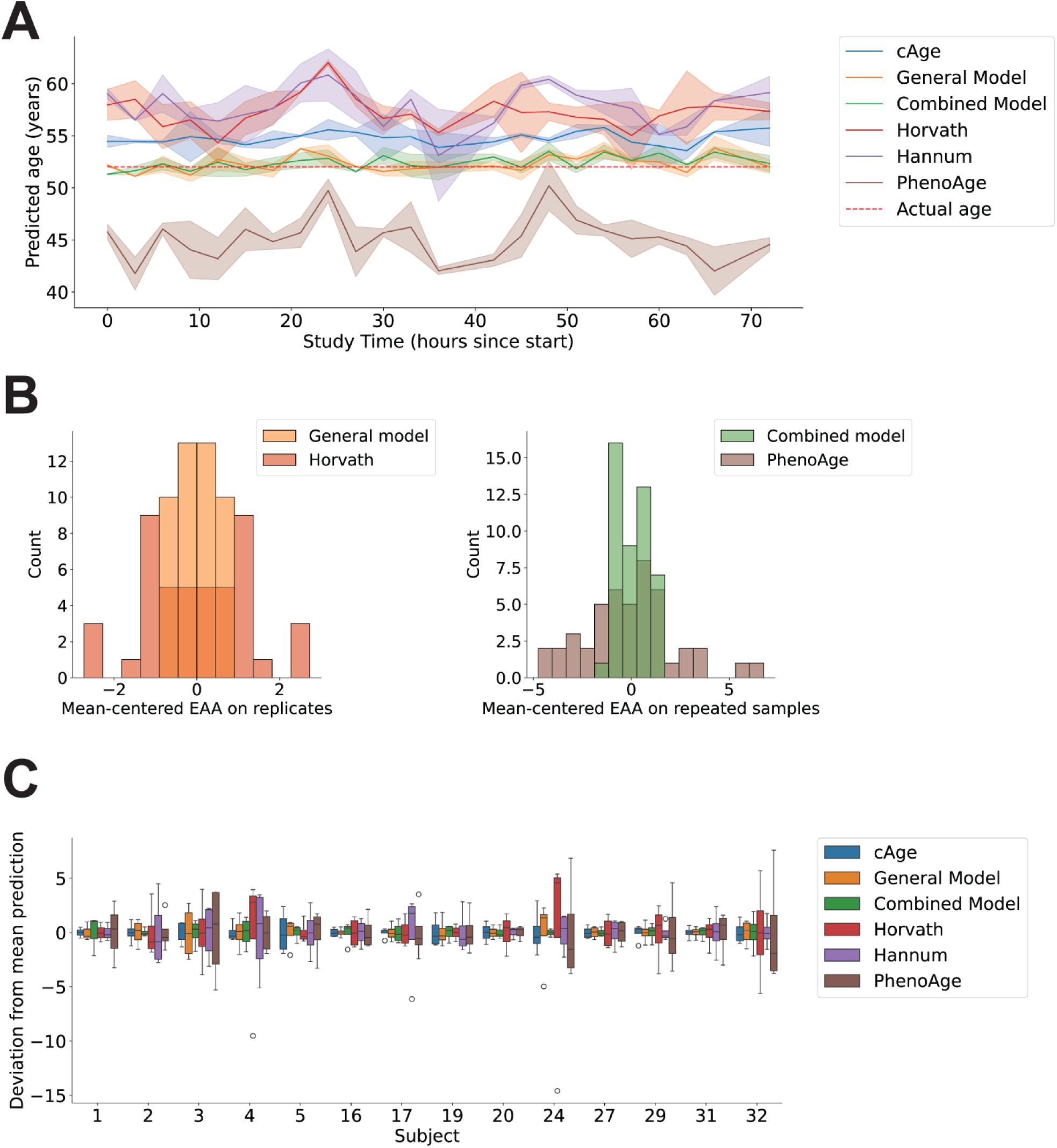
Age predictions technical replicates and repeated samples. **A**. Age values predicted by the different models on the dataset from Koncevičius et al., with multiple samples of the same subject (52-year-old male) obtained at different times and with replicates. Solid lines indicate the mean values, whereas the colored area marks the 95% confidence intervals. **B**. Distribution of mean-centered EAA values on technical replicates (left) and repeated samples from the same subject (right) on the Koncevičius et al. data for the models with the highest and lowest standard deviations. **C.** Distribution of mean-centered age predictions for the 14 subjects from the Apsley et al.’s dataset.

We then applied all the models to the data from Apsley et al., discarding the samples from subjects under the stress test. This data includes DNA methylation data from blood drawn from 31 subjects at 4 different time points over a period of 4 hours and 45 minutes under stress or control conditions^38^ (GSE227809). Using data from the 14 subjects in control condition, we computed the standard deviation across the mean-subtracted predictions of all subjects for all the models. We interpreted this measure as another estimate of σ_subject_. The results were largely compatible with the estimate of σ_subject_ from Koncevičius’ data; the largest difference was observed on Horvath’s model, for which σ_subject_ rose from 1.99 to 3.31 years. In all the other models, σ_subject_ changed by less than 1 year. Also in this dataset, our combined model’s predictions suffered the least variation, whereas those from Horvath’s model changed the most (Figure 5C, Table 1).

According to these results, the sum of technical and intra-subject variation of the EAA predictions represent ∼50 to 60% of the population-wide σ_EAA_ depending on the model. Taken together, the metrics in Table 1 provide a means to judge whether the magnitude of EAA findings puts them beyond the range of normal variations.

### The new epigenetic clock reflects the influence of radiation therapy and breast cancer

Several studies have employed different epigenetic clocks to assess the impact of pathologies and environmental factors on epigenetic age acceleration (EAA)^45^. Using PhenoAge^16^, Qin *et al*. demonstrated that exposure to different anticancer treatments had a significant influence on the EAA of childhood cancer survivors^46^. To validate the performance of our model, we tested it in public datasets that have already shown increased EAA in cancer patients. Using data made public in a subsequent study by Dong *et al*.^47^ (GEO accession number GSE197674) which contained data from 2138 childhood cancer survivors, we studied the influence of radiation therapy (RT) on the EAA determined by cAge and our models. In this dataset, all the models produced epigenetic age predictions highly correlated with chronological age, with r values above 0.9 in all cases (Figure 6A). When comparing subjects who had received RT in one or more body areas (chest, abdomen or pelvis, brain) to those who had not been exposed to RT, all models revealed that the latter group had a significantly lower (p<0.05, one-sided Wilcoxon test) EAA (Figure 6A). This is in line with the results reported in the original study^46^, and similar observations have been made in other studies in which radiation exposure was associated with an increased EAA^48,49^. The rest of the complete and truncated models assign very large EAA values to virtually all samples and do not attribute significantly larger EAAs (p>0.05 in all cases, one-sided Wilcoxon test) to the groups with RT treatments (Supplementary Figure 3). In all cases, the magnitude of these EAA changes is smaller than the σ_EAA_ in the general population (Table 2). Therefore, it is questionable whether the observed EAA differences (here as well as in previous studies) are a true, biologically relevant, reflection of the RT treatment during childhood.

**Figure 6.**
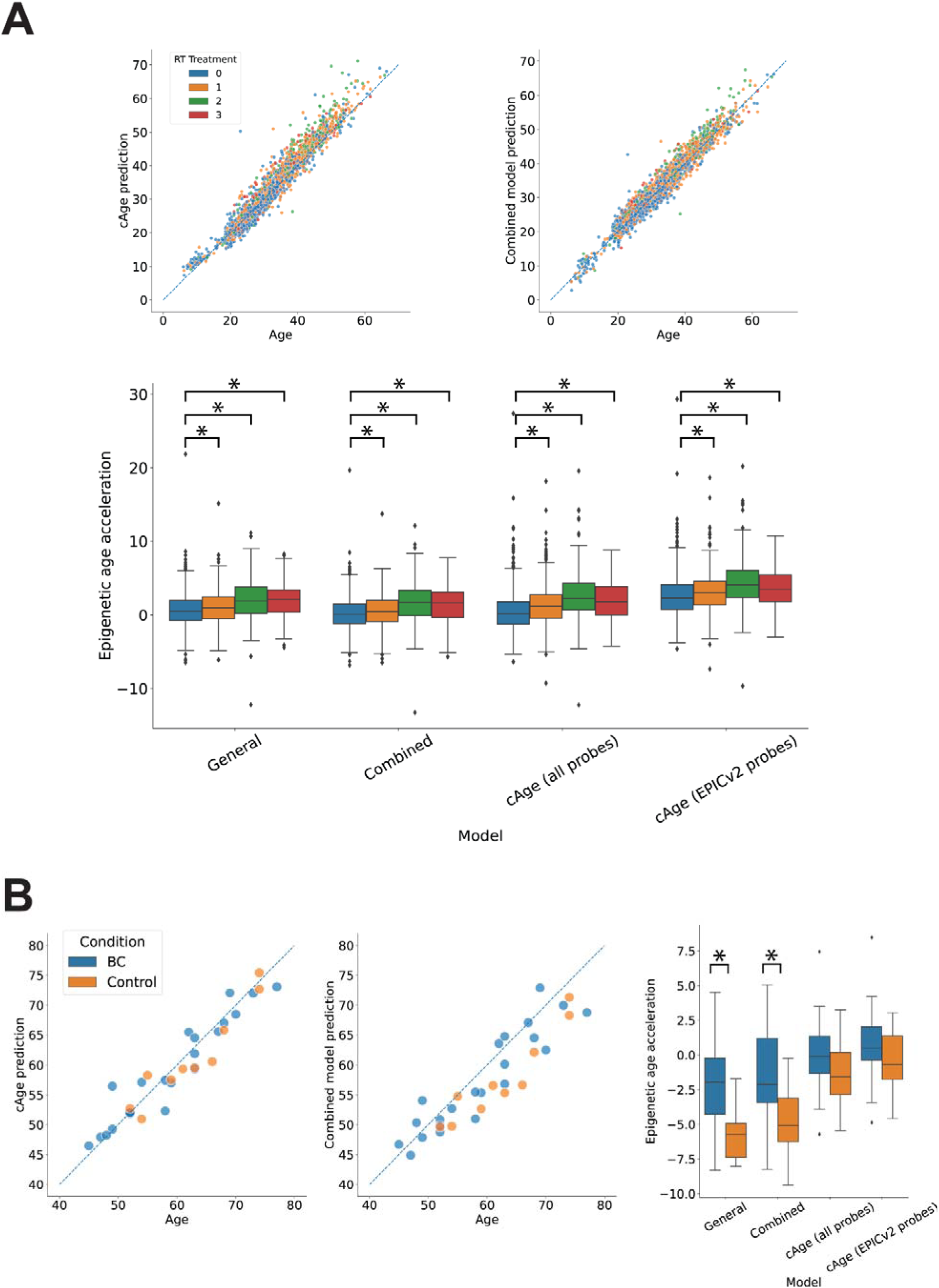
Effects of radiation therapy and spontaneous breast cancer in EAA. **A.** Chronological age and predicted age for childhood cancer survivors, colored by the number of body areas (brain, chest, abdomen/pelvis) in which they received radiation therapy (top). Distribution of the EAA for each group of subjects determined by each of the models (bottom). **B.** Chronological ages of spontaneous breast cancer patients (BC) and control subjects and the corresponding epigenetic ages determined by the combined model and cAge (left). Distribution of the EAA estimated for each group by the different models (right).

**Table 2.** Differences between average EAA of the RT>0 and RT=0 groups observed in the different complete (C) and truncated (T) models.

|  | Hannum<sub>C</sub> | Hannum<sub>T</sub> | Horvath<sub>C</sub> | Horvath<sub>T</sub> | PhenoAge<sub>C</sub> | PhenoAge<sub>T</sub> | cAge<sub>C</sub> | cAge<sub>T</sub> | General Model | Combined Model |
| --- | --- | --- | --- | --- | --- | --- | --- | --- | --- | --- |
| **EAA<sub>RT1-RT0</sub> (years)** | -0,82 | -1,23 | -1,65 | -1,65 | -1,70 | -1,70 | 0,96 | 0,54 | 0,39 | 0,39 |
| **EAA<sub>RT2-RT0</sub> (years)** | -0,19 | -1,07 | -1,68 | -1,68 | -1,87 | -1,87 | 2,37 | 1,73 | 1,34 | 1,48 |
| **EAA<sub>RT3-RT0</sub> (years)** | 0,35 | -0,15 | -0,65 | -0,65 | -0,35 | -0,35 | 2,05 | 1,46 | 1,43 | 1,44 |

Next, we leveraged a dataset with DNA methylation data from peripheral blood leukocytes of sporadic breast cancer patients and control subjects^50^ (GEO accession number GSE148663) to study the effect of the disease on the EAA determined by the different models. We computed the predicted epigenetic age for 22 sporadic breast cancer patients and 10 controls using our models and cAge. Although the correlation between the epigenetic age predictions and the chronological ages of the subjects remained high (0.914-0.951), our models predicted a significantly greater (p<0.05, one-sided Wilcoxon test) EAA in the cancer group (Figure 6B). As reported in the original study^50^, the cancer patients had no previous cancer history and were sampled at the time of diagnosis, so the difference in EAA cannot be attributed to anticancer therapy but rather to the disease itself. The rest of the complete and truncated models do not assign a significantly greater EAA (p>0.05 in all cases, one-sided Wilcoxon test) to the breast cancer group (Supplementary Figure 4). In this case, the magnitude of the EAA differences between cancer patients and controls determined by our models are larger than the σ_EAA_. Therefore, we can consider that these changes do reflect differences larger than the normal variations across individuals in the general population (Table 3).

**Table 3.** Average EAA differences between cancer patients and healthy controls observed in the different complete (C) and truncated (T) models.

|  | Hannum<sub>C</sub> | Hannum<sub>T</sub> | Horvath<sub>C</sub> | Horvath<sub>T</sub> | PhenoAge<sub>C</sub> | PhenoAge<sub>T</sub> | cAge<sub>C</sub> | cAge<sub>T</sub> | General Model | Combined Model |
| --- | --- | --- | --- | --- | --- | --- | --- | --- | --- | --- |
| **EAA<sub>BC</sub>-EAA<sub>Control</sub> (years)** | 2,41 | 2,46 | -0,03 | -0,03 | -0,76 | -0,76 | 1,30 | 1,27 | 3,37 | 3,12 |

Finally, we explored data an interventional study using resistance and aerobic training^51^ (GEO accession number GSE213363). Besides its numerous benefits for overall health^52,53^ and aging^54,55^, physical activity has demonstrated influence on DNA methylation^56,57^. Yet, the study by Furtado et al. reported that after a 16-week exercise intervention, they did not detect significant changes in the epigenetic age calculated by the Horvath model^51^. We applied all the models discussed here to the data from subjects before and after the intervention and found the same result: none of the epiclocks showed significant reductions in epigenetic age after the intervention (p>0.05 in all cases, one-sided Wilcoxon test), neither in the resistance nor in the aerobic training groups (Supplementary Figure 5). This suggests that although the first-generation models examined here can predict chronological age with considerably accuracy, they are less responsive in terms of ability to reflect other biological factors or the effect of certain interventions.

## Discussion

Epigenetic clocks are valuable tools for research on aging and pathological states. In this work, we evaluated the applicability of existing epigenetic clocks to the data generated by EPICv2, the newest methylation microarray model from Illumina. The EPICv2 will phase out previous microarray models (namely, 450k and EPIC), which were used for the development of some widely used epigenetic clocks (e.g., Hannum, Horvath). As the EPICv2 discontinued the use of some of the probes used by existing epiclocks^36^, we felt urged to test whether this would affect the performance of the models.

We generated a large DNA methylation dataset by compiling data from whole blood samples obtained in 24 different studies. This dataset allowed us to quantify the effect of running 4 different epiclocks (Hannum^14^, Horvath^15^, PhenoAge^16^ and the cAge model from Bernabeu *et al*. ^28^) restricted to the set of methylation probes present in the EPICv2 methylation array. Our observations indicate that the results of these 4 models are significantly altered by probes absent from the EPICv2 array.

Epigenetic age models are routinely applied in epigenetic research and are also exploited commercially^58–60^. In the face of commercial discontinuation of the 450k and EPICv1 arrays, our findings suggest that both researchers and commercial vendors alike will need to update their epigenetic clock models to make them compatible with EPICv2 data. Future studies and commercial solutions using this new microarray will require readjusted or new epigenetic age models, which are currently lacking.

As our results show, *none* of the epiclocks tested work as intended on data generated by the EPICv2 microarrays, showing significant differences between the epigenetic ages predicted using data from different chip models. Therefore, we sought to produce a model compatible across the 450k, EPICv1 and EPICv2 microarray platforms, aiming to obtain results in accordance with those of existing methods rather than outperforming them. The approach that we used closely followed the one applied by Bernabeu *et al*. in the development of their cAge model^28^, so that we would rely on established methodologies. Our results on the training and test datasets indicate that our epiclock offered very high performance in the prediction of ages from DNA methylation values. Similar to Bernabeu’s model, these results highlight the benefit of using feature preselection, nonlinear terms, and age-based stratification. We followed a simple strategy regarding these aspects, as we relied on a prior EWAS study^28^ to preselect features, we considered only linear and quadratic features, and we stratified our subjects into three arbitrary age groups. We are confident that follow-up work can improve the results presented here by, for example, conducting more thorough feature selection, deriving more features from the original beta values and/or exploring different data stratification strategies.

Despite its simplicity, our implementation outperformed cAge in the validation dataset, where it obtained significantly lower prediction errors. These results demonstrate that our model can be generalizable (as it is applicable to new data) and that its performance is on par with that of state-of-the-art models.

We tested the variability of epigenetic age predictions on two datasets with repeated samples from the same subjects obtained hours apart from each other. The results indicated that the predictions of cAge and our combined model had the lowest variations across replicates and across repeated samples. We claim that these spontaneous variations of epigenetic age predictions should be considered when examining EAA-related findings: differences smaller than those seen on repeated samples of the same subject obtained on the same day could hardly be considered biologically relevant. Consequently, the significance of previous reports of altered EAAs based on the models d here might need to be re-evaluated.

To demonstrate the ability of our model to reproduce the results of previous models in the detection of EAA alterations, we first applied it to methylation data from cancer survivors^46^. The results of our epiclock indicated an increased EAA induced by radiation therapy, in agreement with previous studies^48,49^. However, the magnitude of this increase (0.39 to 1.44 years) was smaller than the variation we observed in the general population (σ_EAA_). Thus, we cannot claim that the EAA changes detected are reflection of the RT effect on the subjects.

Next, we applied our model to data obtained from breast cancer patients. Our epiclock revealed a significant increase in the EAA of cancer patients compared to that of control subjects, suggesting that the epigenetic age predicted by the model could be sensitive to this particular pathology. In this case, the average EAA difference between the cancer patients and the control group (3.13-3.37 years) was larger than the σ_EAA_ from the non-pathological subjects in the validation dataset (2.43-2.7 years). Therefore, these changes could indeed be a reflection of the pathological state. Notably, the cAge model did not detect a significant difference in the EAA between the control and the cancer groups.

Finally, we analyzed data from an interventional study aimed at improving the health of patients suffering from polycystic ovary syndrome. As in the original study, none of the models we tested reported significant differences in the epigenetic age of the subjects before and after the intervention. Considering the large and extensively documented benefits of physical exercise on health, these results support the idea that despite being good predictors of chronological age, first-generation epigenetic clocks do not necessarily reflect biological factors such as pathological states, environmental exposures or interventions.

As a relevant limitation of our model, we would like to highlight that it is limited to methylation data obtained from whole blood samples, a constraint that is also shared by multiple other models^28,61,62^. Its applicability to data obtained from other tissues has not been assessed. Likewise, we would like to emphasize that its ability to reflect pathological states beyond the sporadic breast cancer cases we have shown here remains to be explored.

Taken together, our results demonstrate that our epigenetic clock is compatible with data generated using the 450k, EPICv1 and EPICv2 microarray platforms. Its epigenetic age predictions are highly correlated with chronological age in control subjects of all ages. Our model exhibited consistency across technical replicates and across repeated samples of the same subjects, with lower variation than the rest of the models tested. As a first-generation epigenetic clock, our model can predict chronological age from DNA methylation data with high accuracy, but its ability to reflect the effects of environmental exposures, pathological states or beneficial interventions is limited. Our work solves a technical barrier derived from technological development that has not yet been addressed and has important implications both for aging research and for biotechnology companies offering services in this field.

## Methods

### Data collection and preprocessing

Methylation data from previous studies were collected from the GEO database^63,64^. In all cases, the data were preprocessed: we used beta values provided by the original authors when available; otherwise, we computed the beta value from the methylated and unmethylated signals using the following formula:

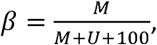

where M and U are the intensities of the methylated and unmethylated signals respectively. The main dataset consisted of 11825 subjects with reported methylation, age and sex data. We discarded 147 with nonnumerical age values, one with a reported age of 891 years, and one with a negative epigenetic age according to the cAge model. To generate training and test sets with even age distributions, we kept only samples with an age value present at least twice in the dataset. This process left a total of 10835 samples on the main dataset, which were split into training and test sets with 7259 and 3576 subjects, respectively.

On the main dataset, we computed the average beta value of each probe across the whole dataset (Supplementary Table 4) and used this average to impute missing values. Among the probes used to train our model (see below) or for running other epigenetic clocks, we found 1.58% missing values. The number of probes missing per dataset is provided in Supplementary Table 5.

The validation dataset included data from 2098 subjects. Excluding subjects with missing or invalid sex and/or age annotations reduced the data to 2095 subjects. A total of 0.23% of the beta values were missing for the set of probes used to run our epiclcok and the cAge models. The missing values were imputed using the average values derived from the main dataset. Supplementary Table 5 reports the number of probes missing per dataset.

The data from studies with repeated samples (GSE247197, GSE227809, 1 and 31 subjects respectively), the cancer survivor dataset (GSE197674, 2138 subjects), the data from the breast cancer study (GSE148663, 32 subjects) and the data from the interventional study (GSE213363, 56 subjects) were also retrieved from the GEO database^63,64^. We used the average beta values derived from the main datasets to impute beta values for 25 missing probes in the cancer survivor dataset.

### Epigenetic age prediction using existing models

We applied the Horvath^15^, Hannum^14^, PhenoAge^16^ and cAge^28^ epiclocks to the methylation data in the main dataset by using the methods and parameters reported by the authors for each case and validated using the implementations in the pyaging Python library^65^. In the case of the Horvath clock, this implies computing an initial value using a linear model and then applying additional transformations for the “young” (<=20) and “adult” (>20) subjects^15^. Similarly, the cAge model uses one model to predict *age* and another to predict log(*age*); if the *age* prediction is <=20, then it is replaced by the exponential of the log(*age*) prediction^28^.

### Training of the new epigenetic clock

We trained a general epigenetic age prediction model using the beta values of 10,000 probes and the squared beta values of 300 probes to perform a regularized linear regression of chronological age using elastic net^40^. The linear and quadratic features were chosen based on an EWAS study^28^: we chose the 10,000 probes with a significant linear association with age with the lowest association p-value and the 300 probes with a significant quadratic association with age with the lowest association p-value.

To train our general epigenetic age prediction model, we separated the main dataset into training and test sets using a 70/30 split stratified by age (i.e., all ages present in the dataset were sampled in both the training and test sets). Elastic net combines L1 and L2 regularizations, which in the scikit-learn implementation^66^ are mixed in proportions given by the parameter L1_ratio, which takes values between 0 and 1. This parameter multiplies the L1 penalty term whereas the L2 term is multiplied by 1-L1_ratio.

Basedon its performance in previous studies, we set the L1_ratio to 0.5^15,67^,. The value of the parameter alpha (i.e. 1/C), which multiplies both the L1 and L2 penalties, was optimized to 0.001 through 5-fold cross-validation.

Age-specific models were trained using the same approach on 3 different batches of data with subjects of specific ages: age<=46 (young), 26<age<69 (middle-aged), and age>=59 (old). The alpha value was set to 0.001 in all cases. The three age-specific models were then used to perform predictions on nonoverlapping *putative* age (age values assigned by the general model) groups: age<=36 (young), 36<age<59 (middle-aged), and age>=59 (old).

## List of abbreviations

EAA: Epigenetic age acceleration
cAge: Chronological age prediction model
EWAS: Epigenome-wide association study
MSE: Mean squared error
RT: Radiation therapy

## Declarations

### Ethics approval and consent to participate

Not applicable.

### Consent for publication

Not applicable.

### Availability of data and materials

The datasets analyzed during the current study are available in the GEO repository. All accession numbers are reported in Supplementary Table 1.

### Competing interests

The authors declare that they have no competing interests.

### Funding

Not applicable.

### Authors’ contributions

L.G. and M.Q. designed the study. L.G. generated and analyzed the data. All authors read and approved the final manuscript.

## Acknowledgements

We are grateful to all the researchers who made their data publicly available and enabled us to carry out this work.

## Notes

### Competing Interest Statement

The authors have declared no competing interest.

### Summary of Updates

We have expanded the scope of the article after peer-review. We have: 1.Modified the title. 2.Added a new set of analyses on the variability of epigenetic age predictions. 3.Included an analysis of data from an interventional study. 4.Added the previously missing information on the statistical tests used; and 5.Revised the text to provide a clearer perspective on the role and capabilities of first-generation epigenetic clocks.

