## Supplementary Figures for "Applicability of epigenetic age models to next-generation methylation arrays"

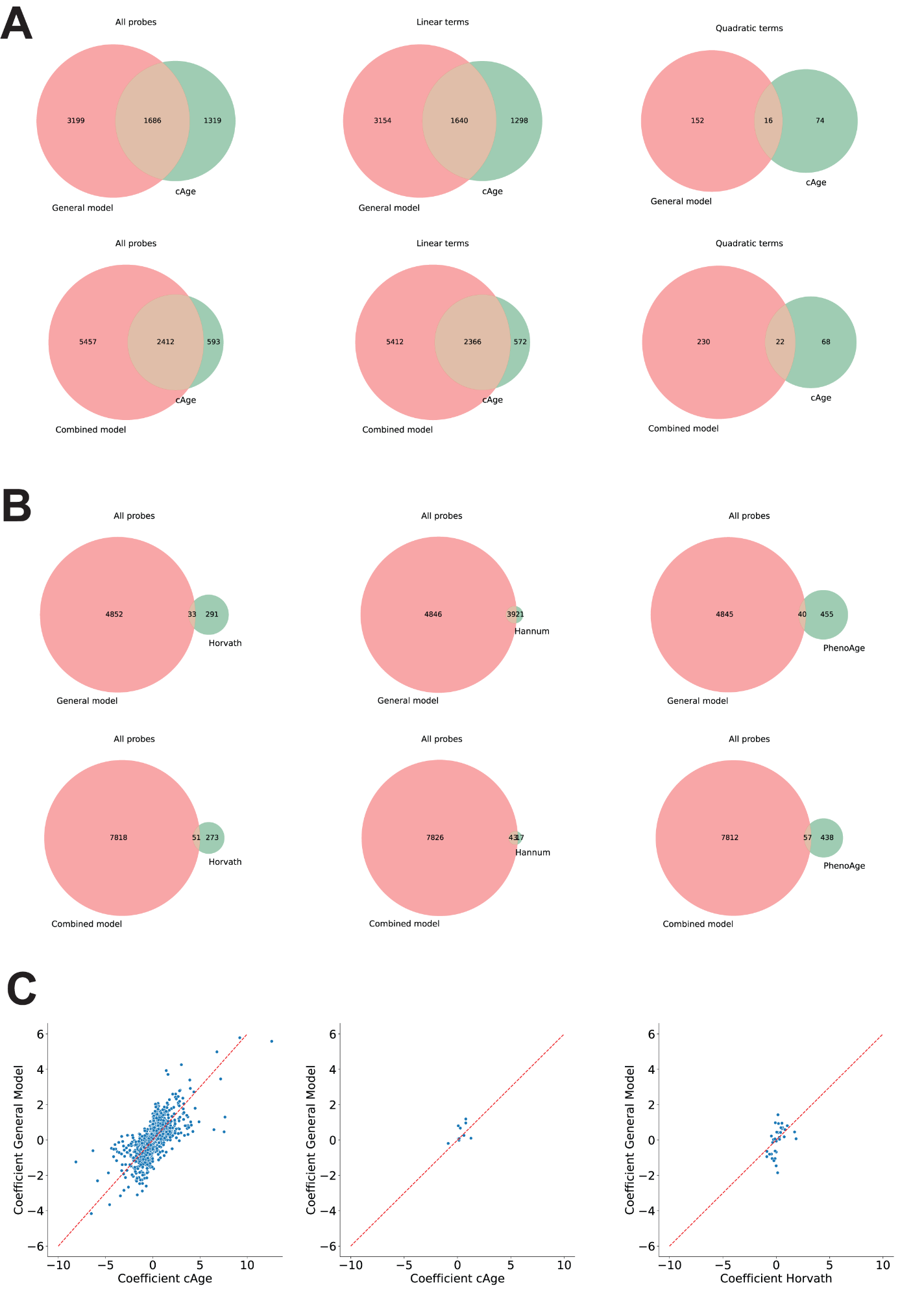

**Supplementary Figure 1 - Overview of differences between the general and combined models and existing epigenetic clocks. A.** Overlap of EPICv2 probes used in the new models and in cAge, considering the full set of probes, only those used in linear terms and only those used in quadratic terms. **B.** Overlap of EPICv2 probes used in the new models and in the Horvath, Hannum and PhenoAge models. **C.** Coefficients used for the EPICv2 probes present in the general model and cAge as linear (left) and quadratic (middle) terms; coefficients used for the EPICv2 overlapping probes in the general and Horvath models, used as linear terms in both cases (right). The red dashed line indicates the position of the x=y identity.

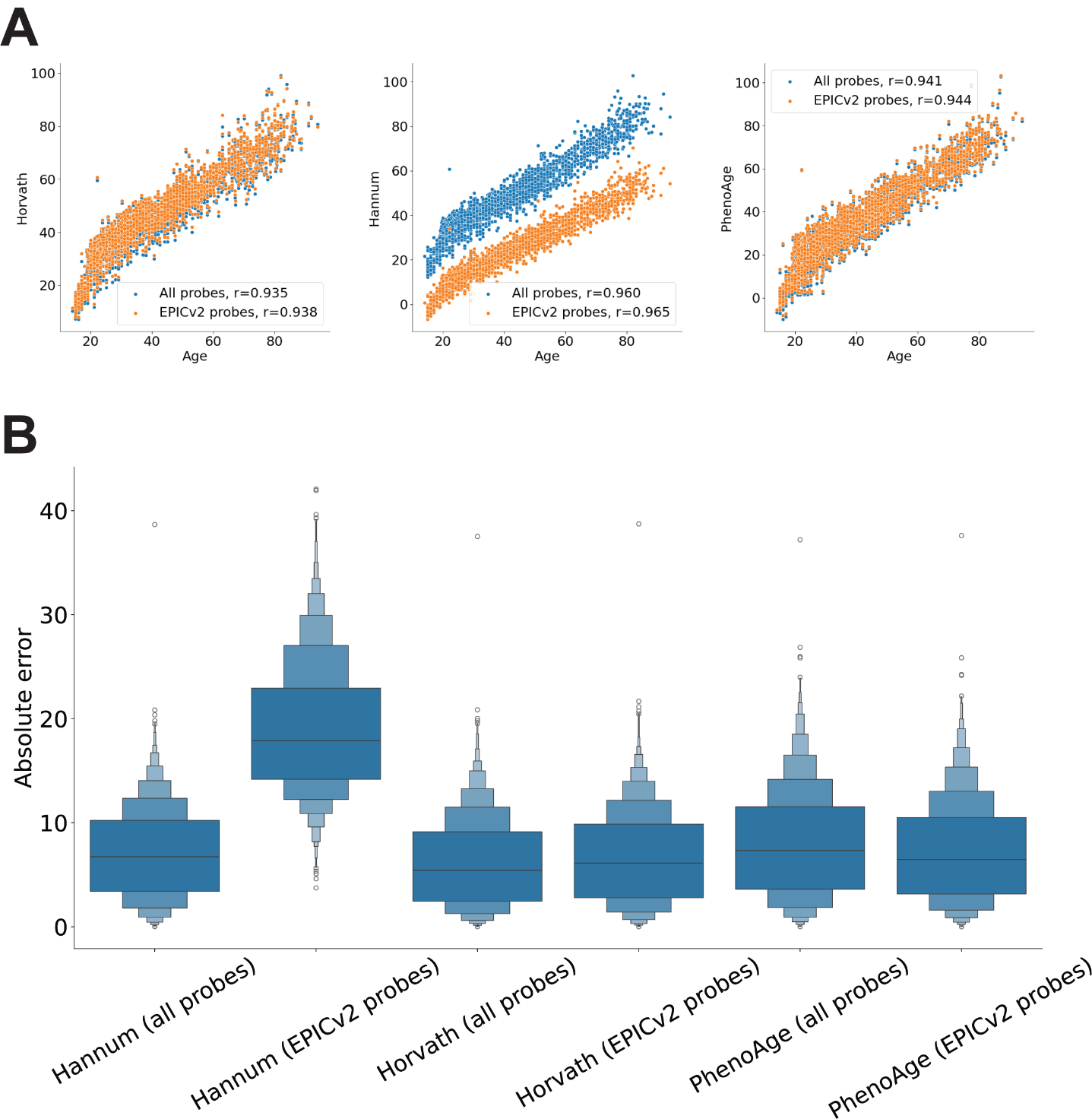

**Supplementary Figure 2 – Predictions of existing models in the validation dataset. A.** Chronological age and predicted epigenetic ages obtained from the Horvath, Hannum and PhenoAge models in the complete (blue) and truncated (orange) forms. **B.** Distribution of absolute error on the age prediction in the validation dataset for the complete and truncated Horvath, Hannum and PhenoAge models.

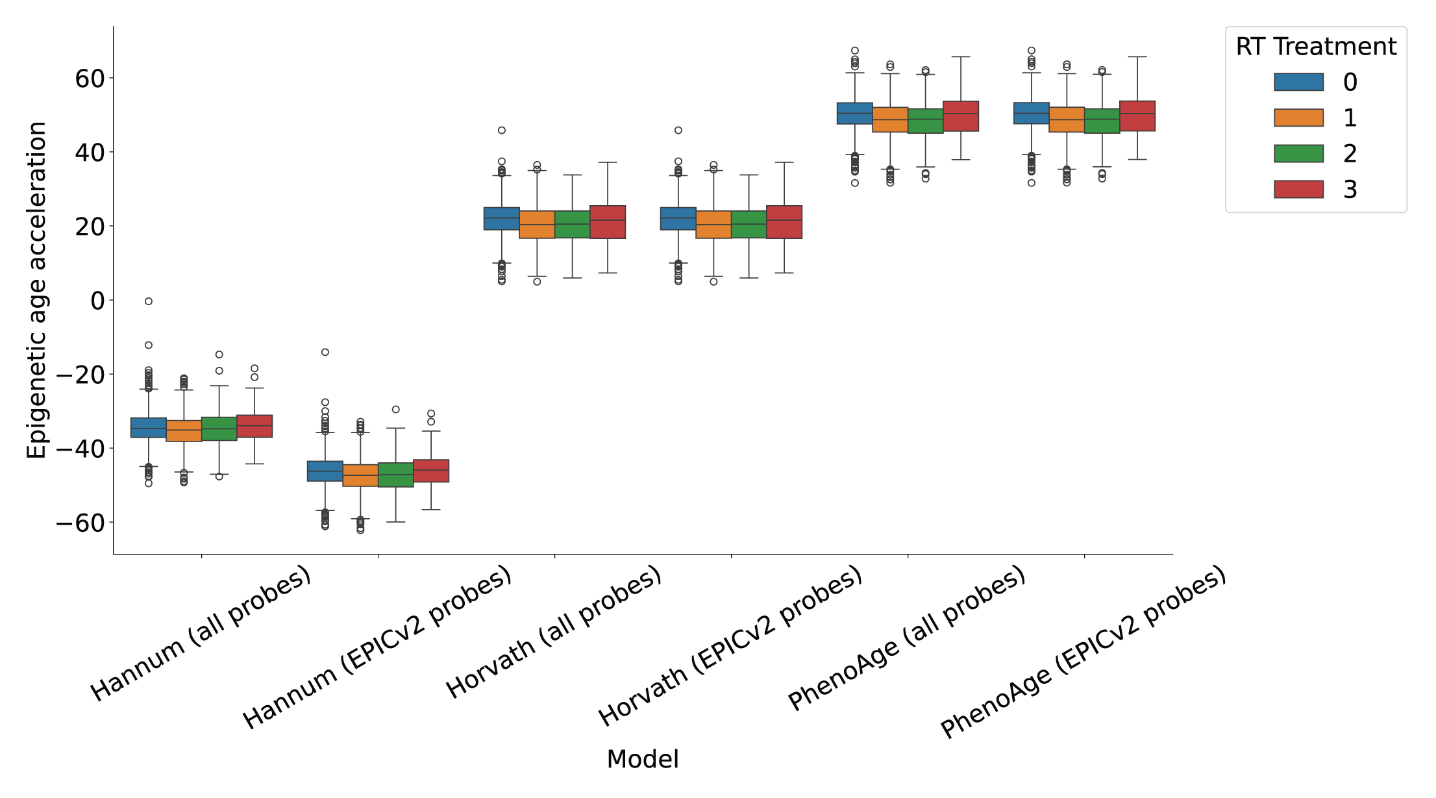

**Supplementary Figure 3 – EAA values in the cancer survivors dataset.** Distribution of EAA values obtained from the complete and truncated Horvath, Hannum and PhenoAge models for samples from subjects with different histories of RT.

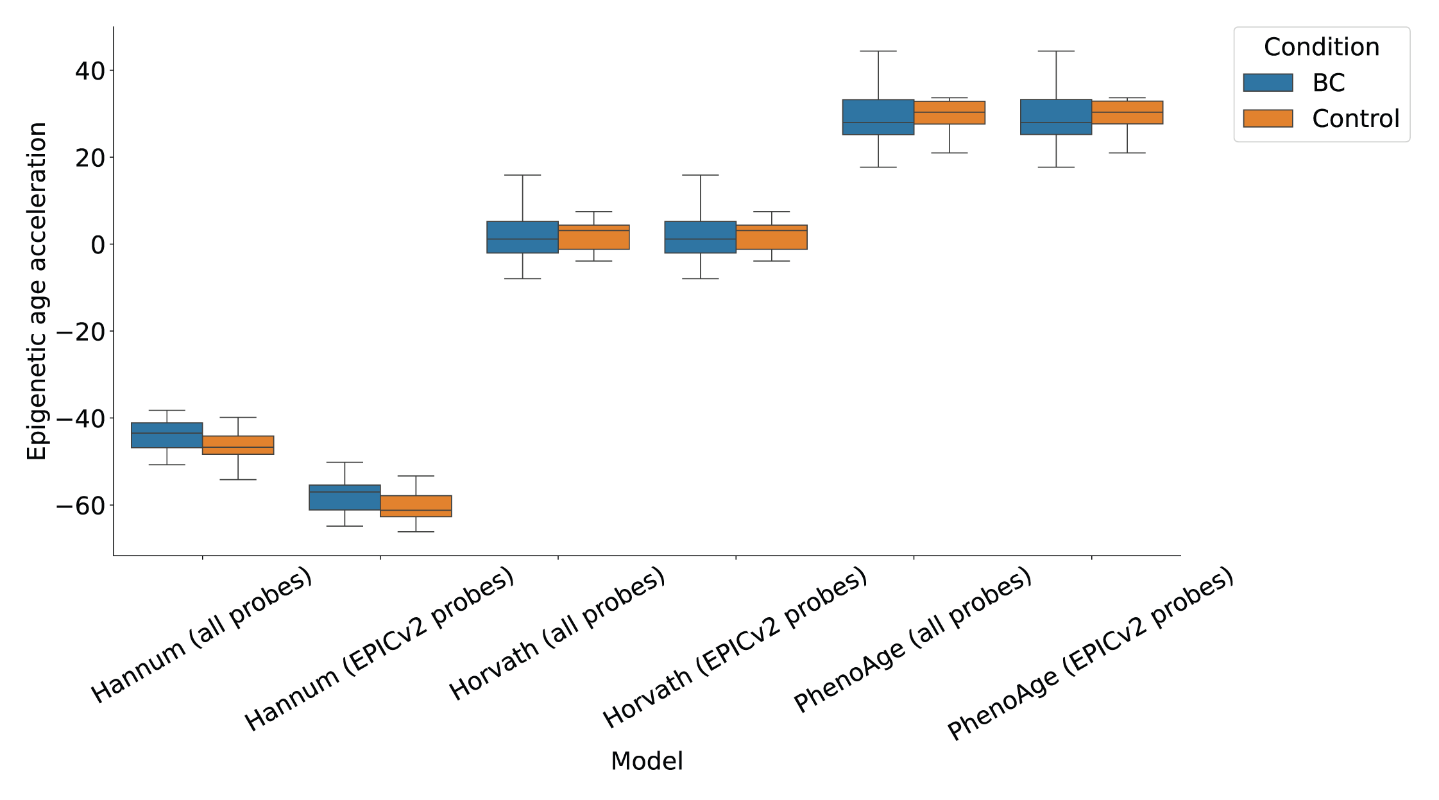

**Supplementary Figure 4 – EAA values in the BC and controls dataset.** Distribution of EAA values obtained from the complete and truncated Horvath, Hannum and PhenoAge models for samples from BC patients (blue) and from healthy controls (orange).

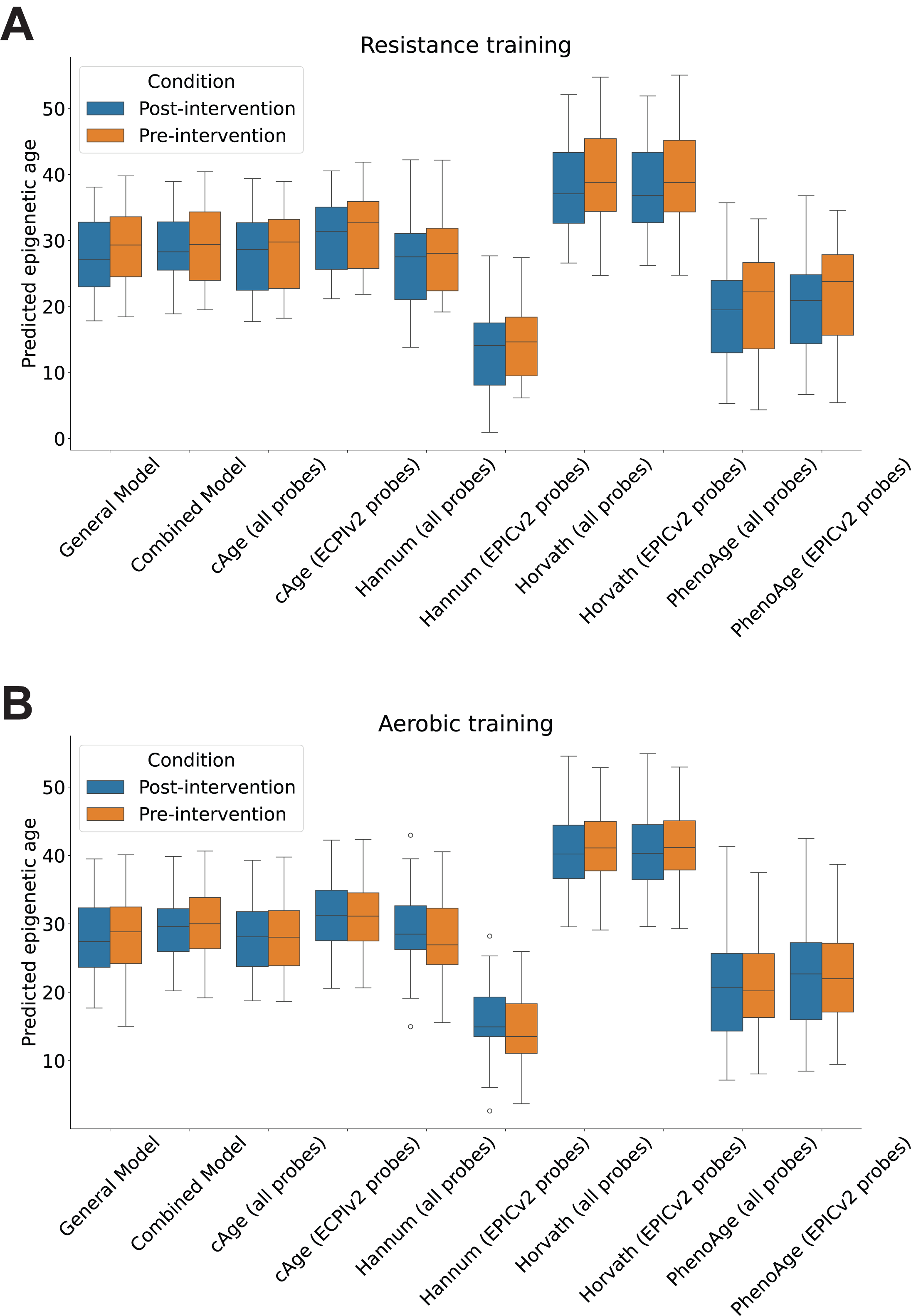

**Supplementary Figure 5 – Epigenetic age predictions in the interventional study data. A.** Distribution of predicted epigenetic ages predicted by each of the complete and truncated models on the data of subjects before and after a 16-week resistance training intervention. **B.** Same distributions for the group of subjects who underwent an aerobic training-based intervention for the same length of time.
